# Exploration of the role of *CHRNA5-A3-B4* genotype in smoking behaviours

**DOI:** 10.1101/818252

**Authors:** Glenda Lassi, Vanessa Tan, Liam Mahedy, Ana Sofia F. Oliveira, Maddy L. Dyer, Katie Drax, Lynne Dawkins, Stephen Rennard, James Matcham, Nicholas J. Timpson, Tim Eisen, Marcus R. Munafò

## Abstract

Genome-wide association studies have identified associations between variation at rs16969968/rs1051730 in the *CHRNA5–A3–B4* gene cluster and smoking related outcomes. Experiments in rodents have described the nicotinic acetylcholine receptors (nAChRs) subunits encoded by this gene cluster and showed a lack of nicotine aversion in nAChRs deficient animal models. We conducted a nicotine challenge and a smoking topography study in humans, hypothesising that: 1. responses to a nicotine challenge would differ according to the rs16969968/rs1051730 genotype and 2. genotype may influence nicotine intake via smoking topography.

We used linear regressions to examine associations between rs16969968/rs1051730 genotype and subjective (questionnaires) and objective (physiological parameters) responses following acute nicotine exposure in never smokers (hypothesis 1) or cigarette smoking in current smokers (hypothesis 2). There was evidence to suggest nicotine exposure increases blood pressure and heart rate, and negatively affects mood, but insufficient evidence that these effects differ by genotype. Carriers of the minor allele following smoking one cigarette, exhibited reduced cravings (b=-2.46, 95% CI -4.87 to - 0.06, p=0.04) and inhaled less smoke per cigarette (b=-0.24, 95% CI - 0.43 to - 0.06, p=0.01) and per puff (b=-0.18, 95% CI -0.32 to -0.01, p=0.02). These results suggest that we need to carefully consider the translational value of the findings of aversion behaviour in nAChRs rodent models, and that deeper inhalation does not explain the strong association between rs16969968/rs1051730 genotype and objective biomarkers of tobacco exposure.

## Introduction

The health, social, and economic consequences of smoking are well known but, despite this, over one billion people continue to smoke worldwide, and each year six million people die prematurely because of tobacco-related illnesses [1]. Reductions in smoking prevalence worldwide have largely been driven by reductions in smoking initiation, while rates of smoking cessation remain low [1, 2]. Cessation is difficult largely because of the addictive nature of nicotine. However, there is considerable inter-individual variability in nicotine dependence and other smoking behaviours. For example, twin studies indicate that genetic factors can explain up to 75% of the variability in smoking behaviour [3, 4].

In 2008, two genome-wide association studies (GWAS) reported that the single nucleotide polymorphism (SNP) rs1051730, in the *CHRNA3* gene, and the strongly correlated SNP rs16969968 in the *CHRNA5* gene in samples of European ethnicity, were associated with nicotine dependence and smoking intensity [5, 6]. Both *CHRNA3* and *CHRNA5* code for nicotinic acetylcholine receptors (nAChRs) subunit proteins, α3 and α5, that, together with the β4 subunit, form the α3(2)α5(1)β4(2) nAChR (see Supplementary Material and Supplementary Figure 1). Circulating nicotine from tobacco smoke binds to nAChRs present in neuronal tissue and the properties of these receptors are determined by the conformation of the subunits. These subunits are characterised by ligand-gated ion channels that open in response to the binding of nicotine and, in doing so, allow the trafficking of cations and provoke a cascade of events at the basis of dependency [7].

**Figure 1.**
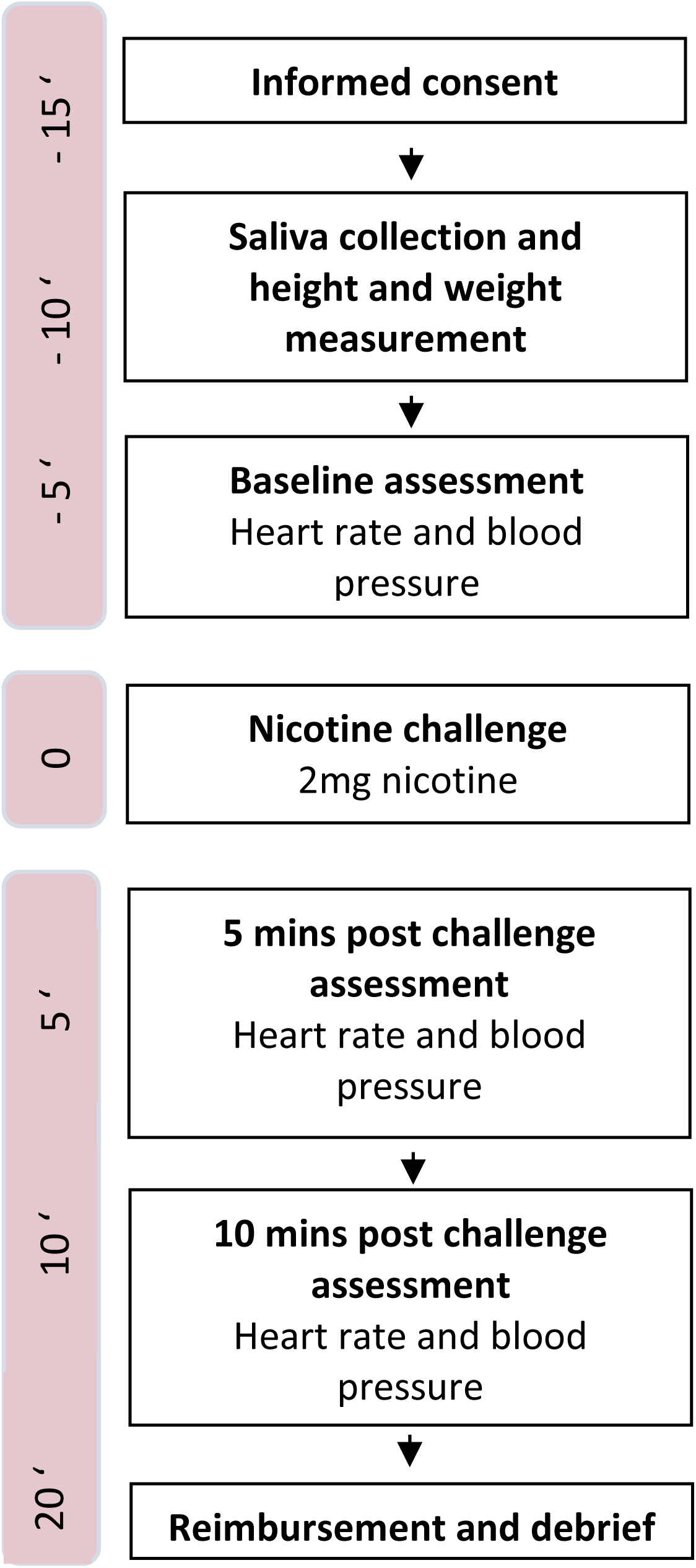
Protocol of the Nicotine Challenge study. Duration of each step of the protocol. The time at which the nicotine challenge was administered was considered time zero (0) and −5’ = 5 minutes prior to nicotine challenge.

These findings have made variants in the *CHRNA5-A3-B4* region promising targets for the study of nicotine dependence, cessation, and smoking intensity. The rs1051730 SNP in *CHRNA3* is a coding, synonymous variant which is in high linkage disequilibrium (LD; R2=1.00 in HapMap 3) with rs16969968. The rs16969968 minor allele is a missense mutation resulting in the substitution of aspartate (D) to asparagine (N) at the 398th amino acid in the resultant α5 subunit protein (D398N) which is likely to reduce the functioning of nicotinic receptors incorporating the α5 subunit (α5*nAChR) [8]. *In vitro* studies have shown that α5*nAChRs need a higher concentration of ACh (the endogenous ligand of nAChRs) to be activated [9] and show a reduced response to nicotine agonists compared with α5 receptor complexes containing the aspartic (major) variant [10].

Animal studies have described the role of the α5 nAChR subunit by investigating the *Chrna5* knockout mouse model’s phenotype. Existing work has shown differential effects of nicotine dose on reward between knockout and wild-type mice using a conditioned place preference task, with knockout mice maintaining place preference at higher nicotine doses [11]. Furthermore, *Chrna5* knockout mice show a more vigorous response to obtain intravenous nicotine infusions. While wild-type mice appeared to titrate the delivery of nicotine dose (through self-administration) to achieve a consistent, desired level, knockout mice did so to a considerably lesser extent, consuming greater amounts as the dose increased [12].

These findings, from *in vitro* and *in vivo* studies, provide insight into the mechanisms through which *CHRNA3/A5* genotype influences smoking intensity and suggest a role for α5*nAChR in mediating this association. A better understanding of how *CHRNA3/A5* genotype influences smoking behaviours could support further investigation of this system as a therapeutic target. Interestingly, pharmaceutical companies, including AstraZeneca [13] and GlaxoSmithKline [14], are of the opinion that clinical development programmes informed by genetically-validated drug targets are less likely to fail [15].

In this study we examined the association of *CHRNA3/A5* genotype with smoking behaviour by studying the response to a nicotine challenge in never smokers and smoking topography in current smokers. Given animal work indicating higher tolerance to nicotine in *Chrna5* knockout animals, we tested objective (heart rate and blood pressure) and subjective (symptoms and mood questionnaires) responses to acute nicotine exposure in never smokers, and whether this differed according to the rs16969968/rs1051730 genotype. By testing never smokers we were able to measure the effects of nicotine while avoiding problems linked with nicotine withdrawal. We also assessed current smokers by recording their smoking behaviours with a portable device to determine the role of smoking topography (*e.g.* volume inhaled when smoking) in the association between the rs16969968/rs1051730 genotype and cotinine, a known smoking biomarker. Various studies indicate stronger associations between *CHRNA5/A3* genotype and cotinine levels rather than self-reported smoking intensity [16, 17], suggesting that rs16969968/rs1051730 variants may influence nicotine intake via other mechanisms than simply number of cigarettes smoked per day.

## Methods

### Nicotine Challenge

#### Participants

Five hundred and thirty-five participants were recruited and tested at two sites: University of Bristol and London South Bank University. All participants were screened upon arrival at the test centre to ensure that they met the study inclusion criteria: i) never-smokers (i.e., consumed fewer than 100 cigarettes over their lifetime) and ii) between 18 and 50 years of age. Participants were excluded if they had used illicit drugs within the previous week, if they were current users of prescription medication, or if they were pregnant or breastfeeding. The study was advertised to the general public via flyers and social media. Recruitment occurred between September 2016 and July 2018.

Participants provided written informed consent before commencing the study, and they were reimbursed for their participation. Ethics approval was obtained from the University of Bristol Faculty of Science Human Research Ethics Committee (#25061523521) and London South Bank University’s Ethical Committee (UREC 1605).

#### Measures

Demographic characteristics were assessed via self-report in the laboratory session. Self-reported sex, ethnicity and age were recorded. Heart rate (HR) (bpm) and blood pressure (BP) (mm/Hg; OMRON Healthcare, UK) were measured at baseline, five and ten minutes after nicotine administration.

Baseline and post-challenge symptoms and mood were measured using the symptoms visual analogue scales (VAS) [18] and the Positive and Negative Affect Schedule (PANAS) [19], respectively. The symptoms VAS is a 10-item questionnaire. Each item is a symptom (such as nausea or dizziness) and the participant indicates how much the symptom is felt, from 0 (not at all) to 10 (extremely). The PANAS comprises two mood scales: one positive and one negative. Each scale consists of 10 items ranging from 1 (not at all) to 5 (extremely).

#### Genotyping

DNA samples were obtained using the Oragene DNA oral collection kit OG-500 (DNA Genotek Inc., Canada) to collect and store saliva. The samples were genotyped for rs1051730 by KASP genotyping assay (LCG Genomics) (see Supplementary Materials for more details).

#### Procedure

The protocol for this study was preregistered in the Open Science Framework (http://doi.org/10.17605/OSF.IO/3G27E). The study comprised a single test session lasting approximately 45 minutes (see Figure 1).

Participants attended either a research centre at the University of Bristol or London South Bank University. Following informed consent, a saliva sample was collected for genotyping purposes, and height and weight were measured. Baseline cardiovascular (i.e., heart rate and blood pressure), VAS and PANAS measures were then completed.

A 2 mg dose of nicotine delivered in mouth spray form (Nicorette® Quickmist) was then administered by the researcher. Cardiovascular and VAS measures were assessed five and ten minutes after nicotine administration. The PANAS was assessed at ten minutes after nicotine administration.

#### Data analysis

We performed paired-sample t-tests to assess the overall effect of nicotine administration on BP, HR, symptoms (VAS) and mood (PANAS). Linear regressions were used to examine the association between rs1051730 genotype (coded additively according to number of copies of the minor allele, T, at this locus) and change in scores (pre-nicotine values subtracted to post-nicotine exposure) for each outcome variable at the five and ten-minute post-challenge time points. The group homozygous for the major allele (CC), previously reported as associated with reduced nicotine dependency and smoking intensity, was the reference. We then assessed the relationship between genotype and change scores on the outcome measures in the total sample, adjusted by ethnicity. In addition, we repeated the analyses only for the European cohort (see Supplementary Table 1 for associations between possible confounders and genotype). All analyses were conducted in Stata version 14.

**Table 1.**
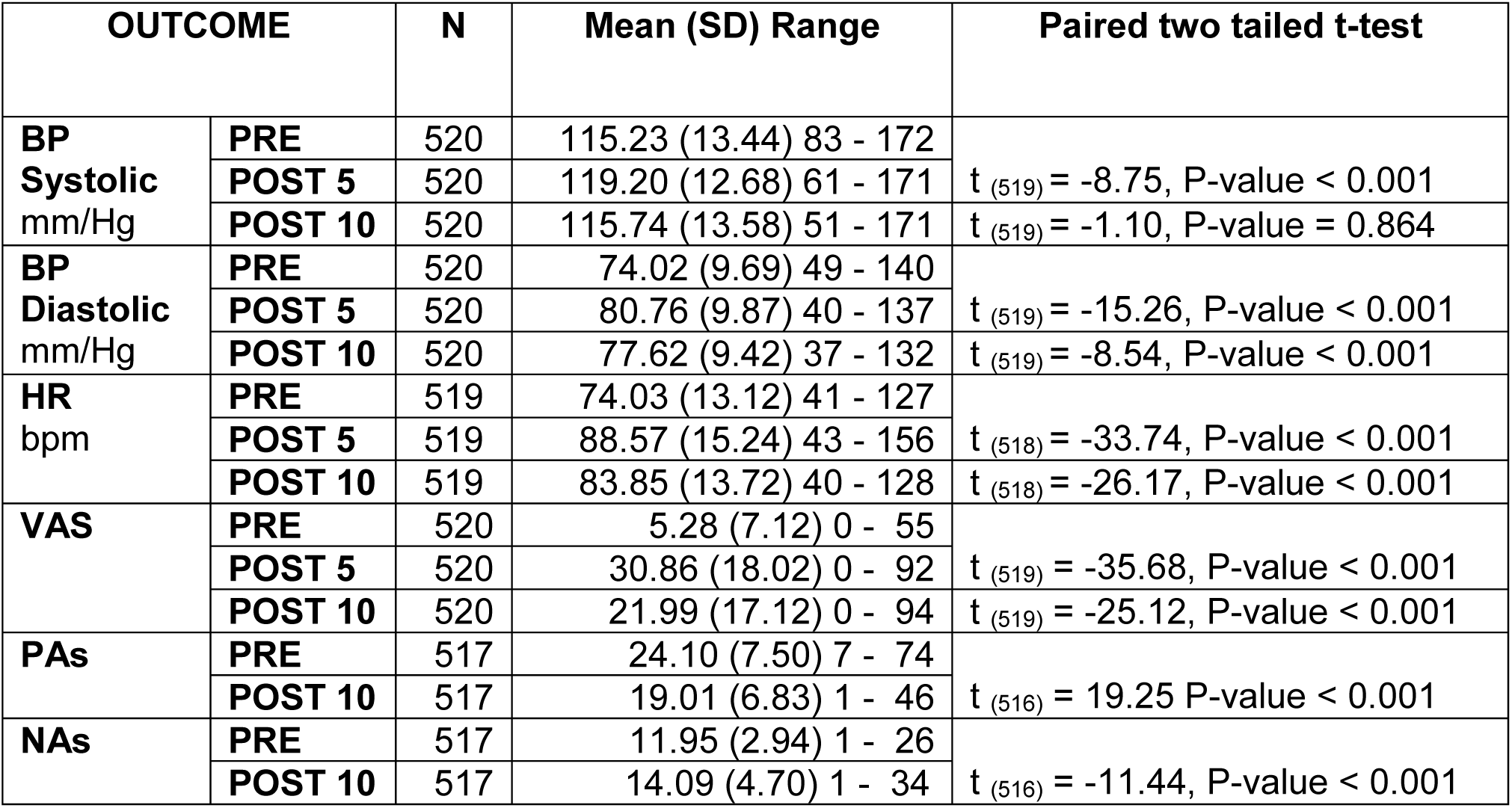
Nicotine Challenge study: cardiovascular and subjective aversion measures. T-tests were performed to assess whether acute nicotine exposure affected heart rate (HR), systolic and diastolic blood pressure (BP), symptoms (measured via the Visual Analogue Scale) and mood (measured via the positive (PAs) and negative affect scale (NAS) at the five and ten-minute post-challenge time points. Negative t-tests results show that the values of the outcomes measured at baseline were smaller than those measured 5 or 10 minutes post nicotine exposure.

### Smoking Topography

#### Participants

All participants were screened for eligibility via a brief telephone interview. Participants were eligible if they were daily smokers (either of hand rolled or manufactured cigarettes) and were in good physical and psychiatric health. Smoking status was confirmed by a carbon monoxide (CO) breath reading (PiCO + Smokerlyzer, Bedfont Scientific, UK) or by a positive urinary cotinine assessment (instant urine test kit, Medical World, UK) when this reading was <10 ppm. Individuals with current substance misuse or dependence (other than nicotine and caffeine), current or past medical or psychiatric illness, and women who were pregnant or breastfeeding were excluded.

Seventy-four of the tested participants were selected from the ALSPAC birth cohort (http://www.bris.ac.uk/alspac) and were recalled-by-genotype (according to homozygosity for the rs16969968/rs1051730 minor or major allele) in an effort to yield an anticipated biological gradient (via genotype). The cohort consists of children born to residents of the former Avon Health Authority area in South West England who had an expected date of delivery between 1 April 1991 and 31 December 1992. The ALSPAC birth cohort consists of 14,541 pregnancies that resulted in 14,062 live births: 13,988 infants were still alive at 1 year [20] and a small number of participants withdrew from the study (N = 24). The sample was further restricted to singletons or first-born twins, resulting in a starting sample of 13,775. Detailed information about ALSPAC is available online www.bris.ac.uk/alspac and in the cohort profiles [20, 21]. The study website contains details of all the data that is available through a fully searchable data dictionary (http://www.bris.ac.uk/alspac/researchers/data-access/data-dictionary/). We then advertised the study with flyers and via social media to the general public and recruited 69 more participants. Recruitment occurred between May 2014 and July 2018. Researchers were blind to the participants’ genotypes since saliva samples were genotyped following the experimental session.

Our target sample size was 200 participants (100 in each of the two genotype groups). Studies of rs1051730 and heaviness of smoking using cigarettes per day indicate a per-allele effect equivalent to approximately one cigarette per day [22]. This corresponds to a 70 ml difference in volume inhaled per cigarette, given an average inhaled volume of 700 ml (SD ∼200 ml), which would be detectable with 70% power (α = 0.05) with this sample size. Studies of rs1051730 and heaviness of smoking using cotinine level indicate a per-allele effect equivalent to a 24.4 ng/ml increase in serum/plasma cotinine level [17]. An effect of this magnitude would be detectable with 80% power (α = 0.05) with our target sample size.

Participants provided written informed consent before commencing the study, and they were reimbursed for their time. Ethics approval was obtained from the ALSPAC Law and Ethics Committee (E2011_02).

#### Genotyping

ALSPAC participants’ genotype was known a priori although the researcher was blind to the genotypes of all participants. The *Genotyping* section of the nicotine challenge study gives details of saliva sample collection for DNA extraction and genotyping of non ALSPAC participants.

#### Procedure

The protocol for the study was published in advance by Ware and colleagues [23] and the procedure is outlined in Figure 2. The study took place over the course of three days. On day one the participant attended the research centre at the University of Bristol for approximately 45 minutes. During this visit the participant provided a saliva sample (to assess cotinine levels) and, if not an ALSPAC participant, a second saliva sample to verify the genotype. The participant was asked to smoke a cigarette using the smoking topography monitor. Cardiovascular, craving, and mood measures were assessed pre- and post-programmed cigarette smoking. Participants were asked to use the smoking topography monitor for each cigarette consumed the day after (day two) in their ‘natural’ environment (i.e., at home). On day three participants returned the monitor.

**Figure 2.**
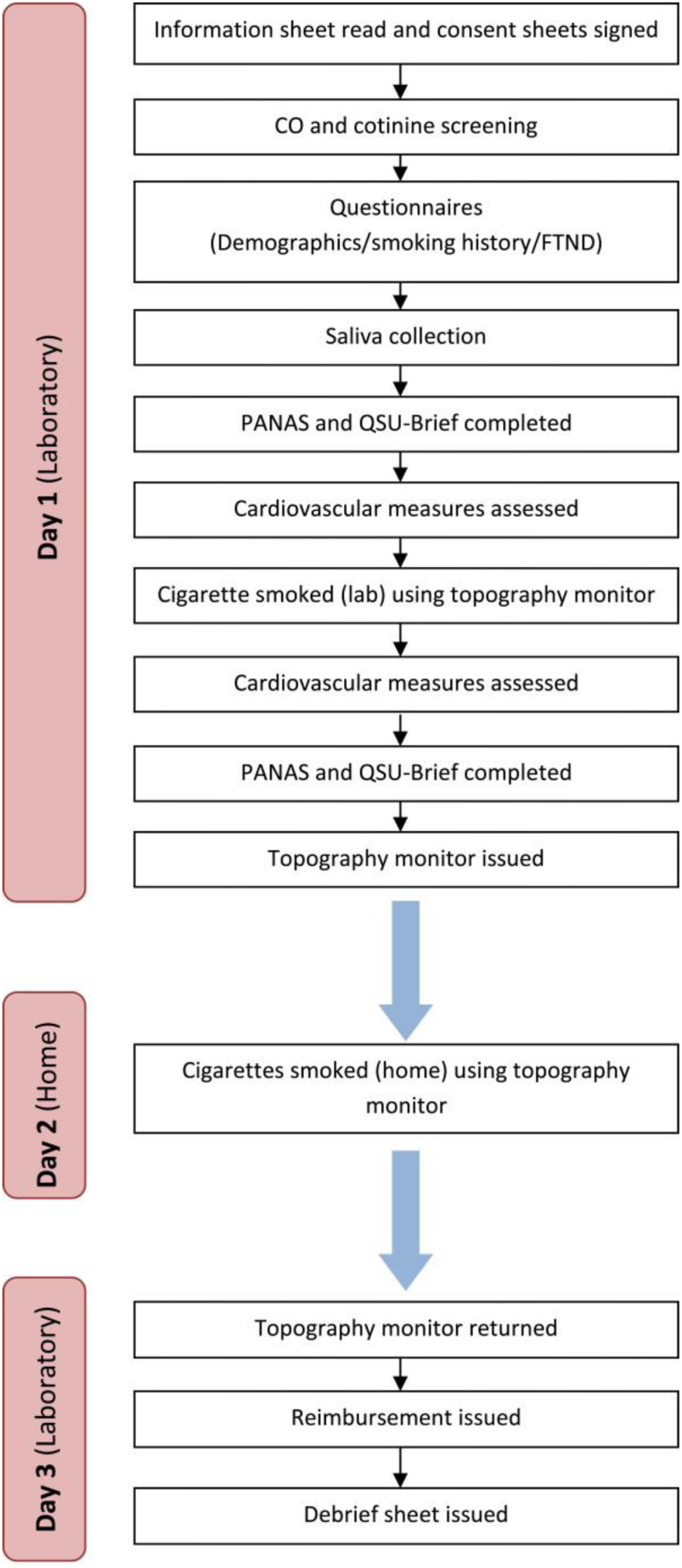
Protocol of the Smoking Topography study. CO = carbon monoxide; FTND = Fagerström Test of Nicotine Dependence; PANAS = Positive and Negative Affect Schedule; QSU = Questionnaire of Smoking Urges.

#### Measures

Demographic characteristics were assessed via self-report in the laboratory session. Self-reported sex, ethnicity and age were recorded. Smoking history was assessed via self-report in the laboratory session, including form of tobacco smoked (manufactured or hand-rolled cigarette) and average cigarettes per day (CPD). CPD data were negatively skewed, therefore CPD was dichotomised (lighter smokers ≤ 10 cigarettes; heavier smokers > 10 cigarettes).

Cotinine levels were assessed from saliva. Saliva samples were collected using salivettes (Sarstedt, Nümbrecht, Germany), centrifuged twice (at 5800 rpm for 15 minutes) within eight hours of collection, frozen (at −30°C) and then sent to ABS Laboratories Ltd which used liquid chromatography with mass spectrometric detection (LC-MS/MS) for the quantification of cotinine content.

Smoking topography was assessed using a smoking topography monitor (CReSS Pocket, Borgwaldt KC, Hamburg, Germany). This is a self-contained, battery-operated device, which captures how a cigarette is smoked as well as temporal measures. Each cigarette was inserted into a plastic mouthpiece that connected to a pressure transducer where inhalation-based changes in pressure were amplified, digitized, and sampled at a rate of 1000 Hz. Smoking topography was assessed both in the laboratory (where the participant smoked one cigarette) and in the participants’ natural environments over the course of one day (all cigarettes smoked from the beginning to the end of the subjective day). Participants were asked to smoke their own cigarettes. We had three main outcome measures for smoking topography. Two of these were measured both in the laboratory and at home and were total volume of tobacco smoke inhaled per puff and per cigarette (ml). The cigarette smoked in the laboratory served the additional purpose to determine the impact of a single cigarette smoked under controlled conditions on cardiovascular, mood and craving measures. Smoking topography measured at home provided a third measure that was the total volume of tobacco smoke consumed per day (ml). Additional smoking topography measures (total number of puffs per cigarette, duration of one cigarette, duration of one puff and inter puff interval) are reported in the supplementary material.

HR (bpm) and diastolic and systolic BP (mm/Hg) were measured, pre- and post-cigarette smoking in the laboratory, using the OMRON M6 blood pressure monitor (OMRON Healthcare, UK).

Smoking craving was assessed via the Brief Questionnaire of Smoking Urges (QSU-Brief) [24] and the PANAS [19] assessed mood. The QSU-brief is a 10-item questionnaire and each item requires a 1 (strongly disagree) to 7 (strongly agree) response. Higher scores indicate a greater urge to smoke. Both questionnaires were administered pre- and post-cigarette smoking.

#### Data analysis

We conducted linear regressions to examine the association of rs1051730 genotype with: a) salivary cotinine level, b) volume inhaled per cigarette, c) volume inhaled per puff, and d) total volume inhaled in a day. Given the non-normal distribution of the variables, cotinine levels were square root transformed and a log transformation was used for all smoking outcomes. In addition, we performed paired two-tailed t-tests to assess the effects of smoking on secondary measures: BP, HR, craving and mood. Linear regression analyses were used to investigate the effect of smoking on these outcomes according to genotype. We tested the association between rs16969968/rs1051730 genotype and the difference between post – and pre-cigarette measures (change scores) of BP (diastolic and systolic), HR, QSU and PANAS scores. We coded rs16969968/rs1051730 genotype additively according to number of copies of minor allele and the participants who were homozygous for the major allele were the reference group.

Sex, and CPD were included as potential confounders of the relationship between rs16969968/rs1051730 genotype and cotinine. Only sex was included in the analyses as potential confounder of the relationship between rs16969968/rs1051730 genotype and cardiovascular and subjective outcomes based on previous literature (see Supplementary Table 3 for associations between confounders and genotype). The results of our analyses are reported fully adjusted. Unadjusted and adjusted analyses are reported in Tables 3-4. Finally, we assessed the correlation between outcomes that were measured both in the laboratory and at home via Pearson correlation coefficient (r). All analyses were conducted in Stata version 14.

Latent profile analysis (LPA) is an extension of latent class analysis (LCA) that can accommodate continuous indicators and was used to examine heterogeneity in patterns of smoking behaviour. LPA models a categorical latent variable that serves to group individuals together who have similar profiles across multiple observed continuous variables. In particular, we selected four variables: i) number of puffs, ii) CPD, iii) smoke inhaled per puff, iv) duration of one cigarette. Starting with a single latent profile, additional profiles were added until model fit were optimised. As there is no common standard for best fit criteria, a number of methods were used. These included: i) information-theoretic methods with lower values indicating better fit to the data i.e., SBIC [25], AIC [26], BIC [27]; ii) likelihood ratio statistical test methods comparing the model with K classes to a model with K-1 classes i.e., LRT [28], bootstrap likelihood ratio test (BLRT) [29], and iii) entropy-based criterion goodness-of-fit indices based on the uncertainty of classification, ranging from 0 to 1 with a high score indicating good fit (entropy) [30]. In estimating our profiles, means were allowed to vary between class, while variances and covariances were constrained to be equal. Associations of risk factors and latent profile membership were examined using the bias-adjusted 3-step analysis as this method accounts for uncertainly in profile assignment Heron and colleagues [31] describe a comparison of approaches to including covariates in latent class models. Analyses were conducted in Mplus 8 [32].

## Results

### Nicotine Challenge

#### Characteristics of participants

Genotype was not available for 15 participants whose saliva sample did not meet the quality standards required for genotyping (N = 520; UoB N = 448 and LSBU N = 72). Overall, 53% of the participants were homozygous for the major allele, 37% were heterozygous and 10% were homozygous for the minor allele. Age ranged from 18 to 50 years (mean age 23, SD 4), 70% of the sample was female, and 77% was of European ethnicity (13% Asian, 3% Black and 7% ‘Other’). The distribution of genotypes by ethnicity and participant characteristics by research site are shown in Figure 3 and values of BP, HR, VAS and PANAS according to research site are shown in Supplementary Table 2. Adjusting for research site did not alter our results substantially, therefore results adjusted for ethnicity only are presented.

**Table 2.** Nicotine Challenge study: association between genotypes and change score. (difference between post – and pre-cigarette measures) of all outcomes at the five and ten-minute post-challenge time points in all participants, and in only the European cohort, regressed on genotype coded additively according to number of copies of minor allele at this locus.

| **ALL**<br>**adjusted by ethnicity** |  |  |  |  |  |
| --- | --- | --- | --- | --- | --- |
| **GENOTYPE** | **Time point** | **N** | **Beta coeff** | **95% CI** | **P-value** |
| **Systolic BP** | POST 5 | 520 | -0.05 | -1.43 to 1.32 | 0.94 |
|  | POST 10 | 520 | -0.49 | -1.88 to 0.91 | 0.49 |
| **Diastolic BP** | POST 5 | 520 | -0.21 | -1.55 to 1.13 | 0.76 |
|  | POST 10 | 520 | -0.75 | -2.03 to 0.53 | 0.25 |
| **HR** | POST 5 | 520 | -0.03 | -1.34 to 1.29 | 0.97 |
|  | POST 10 | 519 | -0.68 | -1.83 to 0.46 | 0.24 |
| **VAS** | POST 5 | 520 | 1.18 | -0.99 to 3.35 | 0.29 |
|  | POST 10 | 518 | 0.07 | -1.95 to 2.09 | 0.94 |
| **PAs** | POST 10 | 519 | -0.04 | -0.87 to 0.79 | 0.92 |
| **NAs** | POST 10 | 519 | -0.08 | -0.48 to 0.64 | 0.78 |
| **EUROPEANS** |  |  |  |  |  |
| **GENOTYPE** | **Time point** | **N** | **Beta coeff** | **95% CI** | **P-value** |
| **Systolic BP** | POST 5 | 400 | -0.22 | -1.67 to 1.90 | 0.75 |
|  | POST 10 | 400 | -0.48 | -1.91 to 0.96 | 0.51 |
| **Diastolic BP** | POST 5 | 400 | -0.41 | -1.83 to 1.01 | 0.57 |
|  | POST 10 | 400 | -0.72 | -1.99 to 0.55 | 0.27 |
| **HR** | POST 5 | 399 | -0.43 | -1.86 to 1.01 | 0.56 |
|  | POST 10 | 399 | -0.86 | -2.10 to 0.40 | 0.18 |
| **VAS** | POST 5 | 400 | 1.04 | -1.22 to 3.30 | 0.37 |
|  | POST 10 | 400 | -0.50 | -2.52 to 1.52 | 0.62 |
| **PAs** | POST 10 | 399 | -0.22 | -1.04 to 0.60 | 0.60 |
| **NAs** | POST 10 | 399 | -0.03 | -0.63 to 0.56 | 0.91 |

HR = heart rate; BP = blood pressure; PAs = positive affect scale and NAs = negative affect scale.

**Figure 3.**
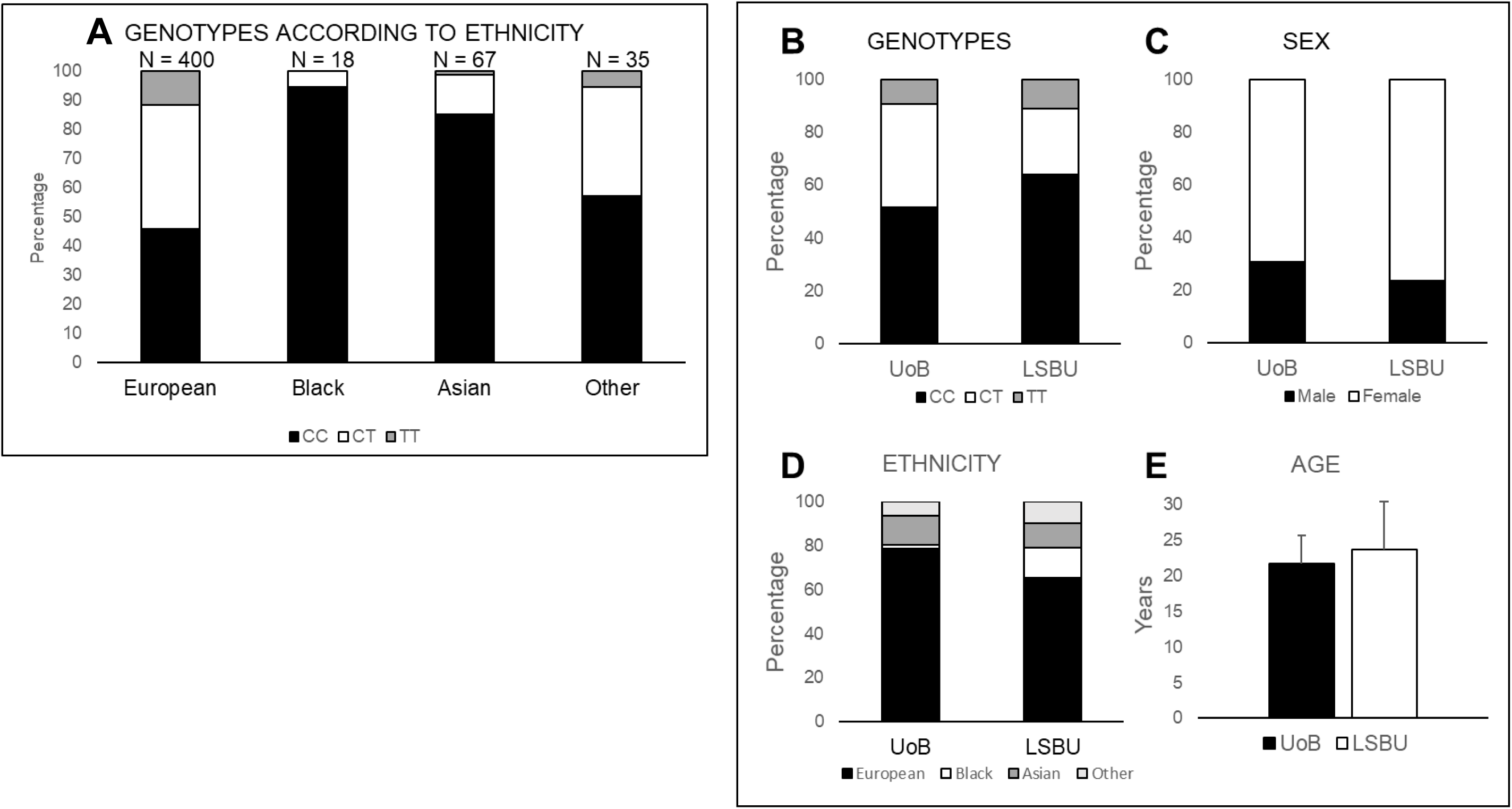
Nicotine Challenge study. Participant characteristics. Participant genotypes according to ethnicity. Here are represented the percentage of participants for each genotype (CC = homozygous for the major allele; CT = heterozygous; TT = homozygous for the minor allele) for each ethnicity (A). Participant characteristics according to research site. Here are represented the characteristics of total sample according to the research site: UoB, N = 448 and LSBU, N = 72, respectively. Percentages of participants for each genotype (B). Percentage of females and males (C). Percentage of participants that were European, Black, Asian and of Other ethnicity (D). Mean age and standard deviation (E).

#### Cardiovascular and subjective response

Table 1 shows the results of paired t-tests assessing the effect of acute nicotine exposure on systolic and diastolic BP, HR, physical symptoms and mood Compared to baseline, systolic and diastolic BP, HR, and VAS (115.23 mm/Hg, 74.02 mm/Hg, 74.03 bpm and score of 5.28, respectively) all increased after 5 minutes (119.20 mm/Hg, 80.76 mm/Hg, 88.57 bpm, and score of 30.96; respectively, all p < 0.001). Ten minutes following the nicotine challenge, diastolic BP, HR, and VAS score were still strongly increased compared to baseline (115.74 mm/Hg, 77.62 mm/Hg, 88.35 bpm and score of 21.99; respectively, all p < 0.001). In addition, negative mood, was 2 points more negative while positive mood was 5 points less positive after nicotine exposure.

#### Associations with genotype

Table 2 shows the results of linear regressions assessing the effect of rs16969968/rs1051730 genotype on cardiovascular and subjective aversion responses to nicotine, captured as change in scores (pre-nicotine values subtracted to post-nicotine exposure) for each outcome variable at the five and ten-minute post-challenge time points. We found no clear evidence of an effect of genotype on change in BP, HR, VAS scores or PANAS scores at 5 or 10 minutes post nicotine challenge. This was evident in both the whole sample and in the European ethnicity subsample.

### Smoking Topography

#### Characteristics of participants

Smoking topography measures were collected in 143 participants. Genotype was not available for 18 ALSPAC participants and two non-ALSPAC participants whose saliva sample did not meet the quality standards required for genotyping. The final sample size was therefore 123 participants (mean age 25, SD 9, years, range 18 to 67 years). Participants were categorized into three groups according to their age: young adult (18 to 24 years; N = 100), adult (25 to 40 years; N = 15), and older adult (47 to 67 years; N = 8). The young adult group was defined according to the age of the ALSPAC participants; ALSPAC young adults’ age at the time of this study was maximum 24 years. We then defined the adult and older adult groups according to the distribution of age and identified a clear age gap of seven years between the two groups. Considering the heterogeneity of these groups (see Supplementary Table 4 for descriptive statistics for Fagerström Test of Nicotine Dependence [FTND] score, cigarettes per day [CPD], and cotinine levels according to age), only the young adult data was analysed.

Fifty-four participants in the young adult group were homozygous for the major allele, 31 were heterozygous and 15 were homozygous for the minor allele. Fifty-one participants were recalled by genotype from the ALSPAC birth cohort and 49 from the general public. All but four participants were of European ethnicity, over 58% of participants were females, 67% used hand-rolled, rather than manufactured, cigarettes, 66% were classified as non-dependent according to their FTND score (< 4) while the remaining 34% scored ≥4, and 64% smoked between 1 and 10 CPD while the rest smoked up to 30 CPD (see Figure 4).

**Figure 4.**
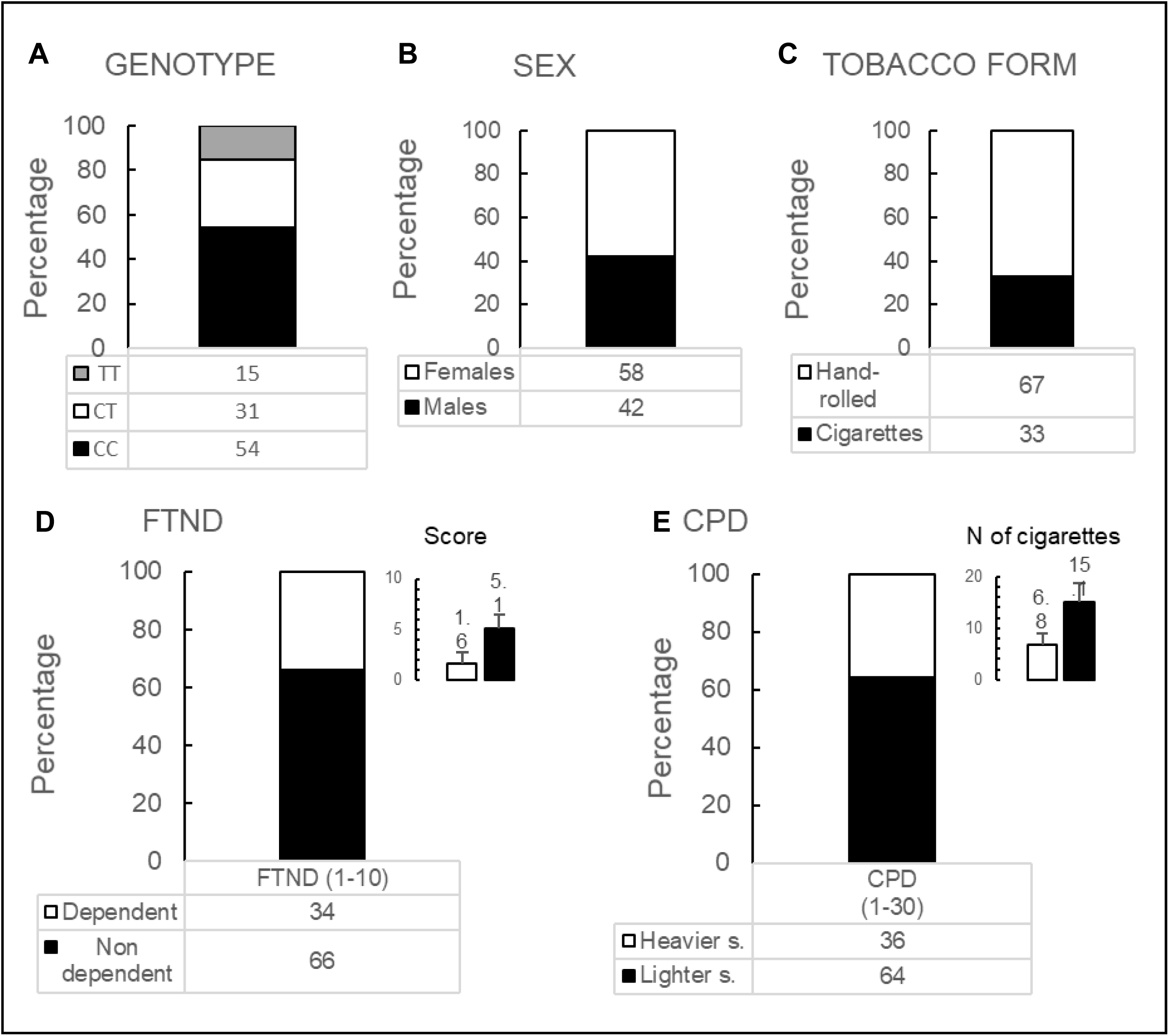
Smoking topography study. Participant characteristics. Percentages of participants for each genotype (CC = homozygous for the major allele; CT = heterozygous; TT = homozygous for the minor allele) in the young people sample (N = 100) (A). Percentage of females and males (B). Percentages of hand-rolled and manufactured cigarette smokers (C). Percentage of dependent (score ≥ 4) and non-dependent (score < 4) smokers and inset representing the average (+SD) FTND score for each category (D). Percentage of light (≤ 10 cigarettes per day) and heavy (> 10 cigarettes per day) smokers with inset representing the average number (+SD) of cigarettes smoked by each category (E). Note that percentages equal the actual figures as the sample N equals 100.

#### Cardiovascular and subjective response

We performed paired t-tests to assess change in a range of outcomes after smoking a cigarette: systolic and diastolic blood pressure (BP), heart rate (HR), smoking craving and positive and negative affect. Both BP (systolic and diastolic) and HR increased after smoking (115.57 mm/Hg, 72.44 mm/Hg, 76.62 bpm at baseline and 119.98 mm/Hg, 76.29 mm/Hg, 88.47 bpm after smoking; all p-values < 0.001), while smoking craving and the positive affect decreased (from 28.86 to 14.33, p-value < 0.001 and from 26.25 to 24.71, p-value = 0.006, respectively; see Supplementary Table 5).

We then analysed the associations between change scores of the cardiovascular, craving, positive and negative affects measures with the rs16969968/rs1051730 genotype. We found little evidence to suggest an association between cardiovascular or mood measures’ change scores and genotype. In contrast, we found some modest evidence of an association between genotype and craving (Table 3). The reduction in smoking craving was greater in carriers of the minor allele (beta −2.46, 95% CI −4.87 to −0.06, p = 0.04).

**Table 3.** Smoking Topography study: association between genotypes and cardiovascular measures and between genotypes and questionnaire measures. Regressions were 1. Unadjusted; or 2. adjusted for sex. PAs = positive affect scale. NAs = negative affect scale. HR = heart rate; BP = blood pressure; QSU = questionnaire of smoking urge.

|  |  | Model 1 |  |  | Model 2 |  |  |
| --- | --- | --- | --- | --- | --- | --- | --- |
| GENOTYPE | N | Beta<br>coeff | 95% CI | P-value | Beta<br>coeff | 95% CI | P-value |
| HR | 96 | -0.39 | -2.86 to 2.08 | 0.75 | -0.30 | -2.77 to 2.17 | 0.81 |
| Systolic BP | 98 | 0.70 | -2.57 to 3.96 | 0.67 | 0.67 | -2.62 to 3.96 | 0.68 |
| Diastolic BP | 98 | -1.04 | -3.77 to 1.67 | 0.44 | -1.05 | -3.79 to 1.68 | 0.45 |
| QSU | 100 | -2.46 | -4.87 to -0.06 | 0.04 |  |  |  |
| PAs | 100 | 0.07 | -1.18 to 1.33 | 0.91 |  |  |  |
| NAs | 100 | 0.26 | -0.52 to 1.05 | 0.51 |  |  |  |

#### Cotinine levels

Mean cotinine levels were equal to 131.13 ng/ml (SD 96.33, range 0.2 to 502.99 ng/ml). The association between genotype and cotinine was positive, indicating that carriers of the minor allele tended to have higher levels of cotinine, but the confidence intervals were wide and there was no clear statistical evidence to support an association (beta 0.27, 95% CI 0.65 to 1.20, p = 0.56; see Table 4).

**Table 4.** Associations of genotypes with cotinine levels and smoking topography outcomes. Model 1 = unadjusted, Model 2 = adjusted for sex, Model 3 = further adjusted for cigarettes smoked per day. Volume of smoke inhaled per cigarette and per puff was measured both in the laboratory (LAB) and at home while the total volume inhaled in 24 hours was measured at home only.

| GENOTYPE | N | MODEL1 |  |  | MODEL2 |  |  | MODEL3 |  |  |
| --- | --- | --- | --- | --- | --- | --- | --- | --- | --- | --- |
|  |  | Beta coeff | 95% CI | P-value | Beta coeff | 95% CI | P-value | Beta coeff | 95% CI | P-value |
| **COTININE levels** | 98 | 0.59 | - 0.54 to 1.74 | 0.30 | 0.58 | -0.56 to 1.72 | 0.32 | 0.27 | -0.65 to 1.20 | 0.56 |
| **LAB**<br>**Volume inhaled from**<br>**one cigarette** | 92 | -0.24 | -0.43 to -0.06 | 0.01 | -0.24 | -0.43 to -0.06 | 0.01 | -0.25 | -0.43 to -0.06 | 0.01 |
| **LAB**<br>**Mean volume of**<br>**smoke inhaled per**<br>**puff** | 92 | -0.17 | -0.32 to -0.01 | 0.03 | -0.17 | -0.32 to -0.02 | 0.03 | -0.18 | -0.33 to -0.03 | 0.02 |
| **HOME**<br>**Mean volume inhaled**<br>**from one cigarette** | 94 | -0.10 | -0.24 to 0.05 | 0.19 | -0.10 | -0.25 to 0.04 | 0.18 | -0.10 | -0.25 to 0.04 | 0.16 |
| **HOME**<br>**Mean volume of**<br>**smoke inhaled per**<br>**puff** | 94 | -0.12 | -0.26 to 0.01 | 0.08 | -0.13 | -0.27 to 0.01 | 0.07 | -0.13 | -0.27 to 0.01 | 0.06 |
| **HOME**<br>**Total volume inhaled**<br>**in 24 hours** | 92 | -0.05 | -0.27 to 0.17 | 0.65 | -0.06 | -0.28 to 0.16 | 0.59 | -0.11 | -0.28 to 0.06 | 0.22 |

#### Smoking topography

For the data collected in the laboratory, there was some evidence for a negative association between genotype and volume of smoke inhaled per cigarette (beta −0.24, 95% CI - 0.43 to - 0.06, p = 0.01) and per puff (beta −0.18, 95% CI - 0.32 to −0.01, p = 0.02; see Table 2 and Figure 5A), with lower volume inhaled in carriers of the rs16969968/rs1051730 minor allele, known to be positively associated with smoking intensity.

**Figure 5.**
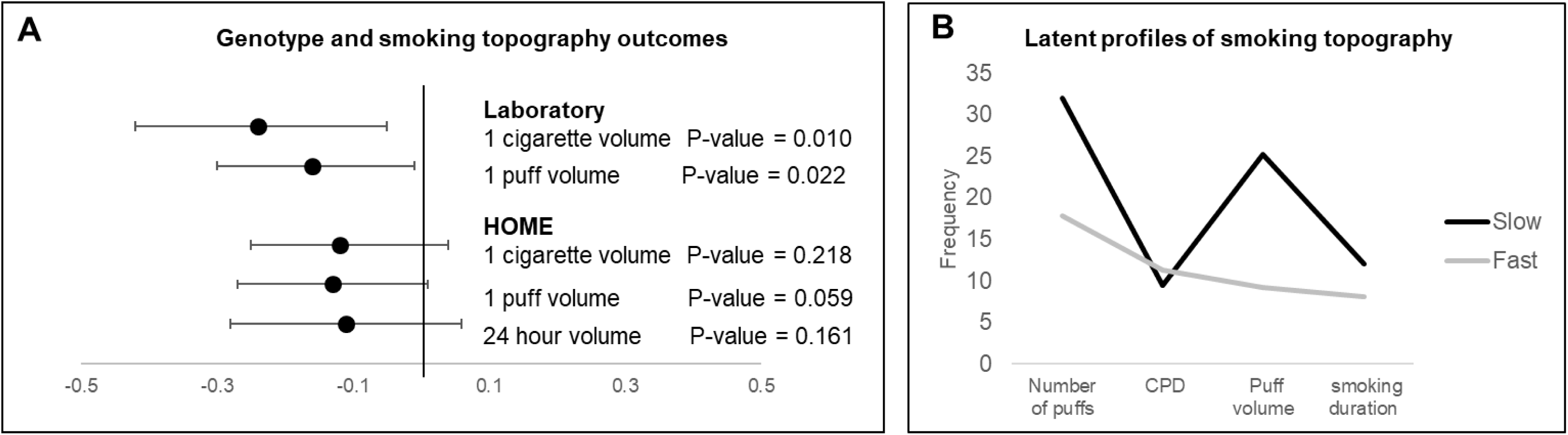
Associations between genotype and smoking topography outcomes. Dots represent the beta coefficient, horizontal lines represent 95% confidence intervals. Associations between genotype and smoking topography outcomes measured in the laboratory: volume inhaled smoking one cigarette and mean volume inhaled per puff, and at home: mean volume inhaled per cigarette, per puff and over 24 hours. CC = homozygous for the major allele; CT = heterozygous; TT = homozygous for the minor allele (A). **Latent profile analysis (LPA).** The LPA identified two smoking patterns. One smoking pattern, the ‘fast smokers’, is characterised by fewer puffs, more cigarettes smoked per day (CPD), and more smoke inhaled per puff, compared to the other smoking pattern, the ‘slow’ smokers, that presents with the opposite pattern. N = 140. Puff volume is represented in deciliters (dl) and smoking duration of one cigarette is represented in units of 30s (B).

For data collected at home, and in contrast with the laboratory measures, there was weak evidence of an association between genotype and volume of smoke inhaled per cigarette (beta −0.10, 95% CI −0.25 to - 0.04, p = 0.22) and per puff (beta −0.13, 95% CI −0.27 to −0.01, p = 0.06), although the direction of the association remained negative. There was insufficient evidence of an association between genotype and total volume of smoke inhaled in 24 hours (beta −0.11, 95% CI −0.28 to 0.06, p = 0.16; see Table 4 and Figure 5A). Summary statistics for smoking topography outcomes are shown in Supplementary Table 6.

The outcome measures in the laboratory were correlated with those assessed at home: r = 0.72 for the volume of smoke inhaled per cigarette, r = 0.78 for the volume of smoke inhaled per puff, r = 0.73 for CPD self-reported and CPD as measured by the smoking device (p < 0.001; see Supplementary Table 7).

We found no clear evidence for an association between the rs16969968/rs1051730 genotype and additional smoking topography outcomes (number of puffs per cigarette, cigarette duration, puff duration and inter puff interval; see Supplementary Tables 8 – 9).

#### Patterns of smoking behaviour

To test whether the smoking topography data presented with smoking patterns, we ran a latent profile analysis. A two-class model was adequate to explain the heterogeneity in the smoking behaviour data based on non-significant Lo-Mendell-Rubin likelihood ratio test (LRT) and bootstrap likelihood ratio test (BLRT) values when using three classes (indicating that the previous class was the best fitting class). There was a minimal decrease in Akaike’s Information Criteria (AIC), Bayesian Information Criterion (BIC), and sample size adjusted Bayesian Information Criterion (SBIC) between the two- and three-class models, further supporting selection of the two-class model (Supplementary Table 10). The two-class model comprised patterns of smoking behaviour that were labelled as ‘Fast’ (69%) and ‘Slow’ (31%) (Figure 5B). The ‘fast smokers’ took a smaller number of puffs per day, smoked more cigarettes per day, and inhaled more smoke per puff, compared to the ‘slow’ smokers.

To assess the association between genotype (homozygous for the major allele and carriers of the minor allele) and smoking behaviour profiles, coherently with the main analyses, we restricted the analysis to participants with available information on genotype exposure and were aged 24 years and under (N = 100). The association was negative, indicating that carriers of the rs16969968/rs1051730 minor allele were less likely to belong to the slow smokers’ category, but confidence intervals were wide and there was no clear statistical evidence to support this association (beta −0.95, 95% CI −2.20 to 0.29; p = 0.13).

## Discussion

We tested the associations of rs16969968/rs1051730 genotype with subjective and objective responses to acute nicotine exposure in never smokers and cigarette smoking in current smokers. We found evidence to suggest nicotine exposure and smoking increase systolic and diastolic BP and HR and they negatively affect mood. Our findings suggest that individuals experience an aversive response to acute nicotine exposure, but we found little evidence to suggest that this is associated with *CHRNA5/A3* genotype. In addition, we found that following smoking one cigarette, craving reduction is more evident in carriers of the rs1051730 minor allele, reported in GWAS as associated with greater consumption of cigarettes and nicotine dependence, and that these individuals inhale less smoke both per cigarette and per puff.

### Comparison with previous studies

In the nicotine challenge study, we found that systolic and diastolic BP, HR, and VAS symptom score were higher five minutes after an acute exposure to nicotine compared to baseline, and this increase persisted up to ten minutes. In a similar way, negative mood increased while positive mood decreased. However, we found insufficient evidence of any differences by rs16969968/rs1051730 genotype. Animal studies have shown that *Chrna5* knockout mice exhibit an increase in self-administration of high nicotine doses compared to wild type mice [12, 33], at least in part due to an attenuated aversive response to nicotine; however, the data reported here does not support this hypothesis in humans. While the animal work has led to crucial insights into the role of the a5 subunit in the reward circuitry [34], there are important differences between mice and humans. First, *Chrna5* knockout mouse models are used because mice do not carry orthologous variants to rs16969968/rs1051730, although humanized models have now been engineered [35]. Second, nicotine pharmacokinetic profile and plasma levels after nicotine administration vary and are difficult to compare across species [36]. One study that focused on aversion behaviour and rs16969968 genotype in humans used a sample of smokers who were kept abstinent for one night and intravenously infused with nicotine. Withdrawal, urge to smoke (QSU), mood (PANAS), aversiveness, HR and BP were assessed at baseline and subsequently after IV nicotine infusions, and ratings of withdrawal, urge to smoke and aversive effects in response to nicotine were lower for minor allele carriers [37]. Critically, this study examined responses in abstinent smokers, hence modelling the smoking relapse, whereas we examined responses in never smokers, hence modelling smoking initiation.

In our smoking topography study, smoking a cigarette also increased HR and BP (both diastolic and systolic). In addition, smoking reduced positive affect but did not influence negative affect. All these effects did not appear to differ substantially by genotype. Interestingly, cigarette craving declined after smoking and minor allele carriers exhibited a greater reduction. This is consistent with the findings of Jensen and collegues [37], that investigated the aversive effect of nicotine in abstinent smokers, and which also reported reduced craving in carriers of the rs16969968 minor allele but no differences in heart rate and blood pressure by genotype. The association between the rs16969968/rs1051730 genotype and cotinine levels was in the expected direction: carriers of the minor allele showed higher cotinine levels, although the analysis was underpowered to detect differences in cotinine levels according to genotype. Unexpectedly, we found a negative association between genotype and volume inhaled (both per cigarette and per puff) for smoking topography measures collected in the laboratory. Minor allele carriers inhaled less volume of smoke per puff (and did not take more puffs, see Supplementary Table 9), and this was robust to adjustment for CPD. Furthermore, we classified the complete dataset according to two latent patterns: ‘slow’ and ‘fast’ smokers. When we assessed whether genotype predicted membership to one of the two classes, we found very weak evidence that carriers of the minor allele were less likely to be slow smokers and inhaled less smoke per puff, further confirming our finding on the negative association between volume of smoke inhaled and the rs16969968/rs1051730 genotype.

Our hypothesised mechanism that rs16969968/rs1051730 variants may influence nicotine intake via other mechanisms than just CPD (e.g., via deeper inhalation) was not supported with the data collected here. This hypothesis was based on the results of previous work [16, 17] which showed that the association between cotinine levels and rs16969968/rs1051730 was stronger than the previous reported associations between CPD or FTND and rs16969968/rs1051730 genotype. This relationship between cotinine and genotype almost did not change when adjusting for CPD [17]. If the same number of CPD can lead to different cotinine levels, we hypothesised that this was due to differences in smoking topography. Smoking topography measures are unavailable for large GWAS because they are resource demanding, hence we conducted this study to assess the relationship of smoking topography outcomes with the rs16969968/rs1051730 genotype.

Importantly, we recruited a young sample, with mean age ∼21 years and range 18 to 24 years, compared to the samples used in previous studies that looked at the association between rs16969968/rs1051730 genotype and cotinine or CPD [6, 10, 17, 38-40]. Therefore, the smokers tested in this study were young people, still relatively light smokers, who may not have achieved full dependence, with genotype possibly having weaker effects on phenotype as a result.

Our findings were stronger for the tests conducted in the laboratory. Tests conducted on data collected at home exhibited the same direction of association with genotype as the laboratory measures, despite our home topography estimates being imprecise and the statistical evidence for association weak. The smoking device used to collect the smoking topography data was calibrated once in the laboratory for the cigarette that was carefully inserted by the researcher. Subsequent cigarettes, smoked at home, were inserted in the device by the participant and this may have added some noise to the values that were averaged (over the total number of cigarettes and puffs) and used for the analyses.

### Limitations

There are a number of limitations that should be considered when interpreting results from both studies here. First (and relating to nicotine challenge), smoking status was self-reported and not biochemically verified therefore can be inaccurate; yet previous studies have shown high concordance between biomonitoring of smoking and self-reported smoking [41, 42]. Second, data were collected across two research sites, which may have introduced noise into our measures. However, we repeated analyses adjusting for research site and the results were not substantially different. Third (and relating to smoking topography), we were unable to reach our target sample size, meaning that the study was likely underpowered. Fourth, we required participants to use smoking topography monitors outside of the laboratory, and this is likely to have introduced noise to our measures; for this reason, we have focussed on the measures collected in the laboratory and then compared them to those collected at home. Fifth, it is possible that smoking devices may affect smoking behaviour. For example, Ross and Juliano [43] found that using these devices reduced the rewarding effects of smoking; nevertheless this would have probably equally affected all participants. Last, with this study we cannot exclude that our hypothesised mechanism may be also influenced by other genes such as, for example, *CYP2A6* that, together with the *CHRNA5-A3-B4* gene cluster, is, to date, among the strongest candidate therapeutic targets for smoking treatment.

### Conclusion and Implications

Our findings suggest that we need to carefully consider the translational value of findings that support lack of aversion behaviour in nAChRs rodent models. Moreover, results relating to smoking topography did not provide evidence to support our hypothesis that deeper inhalation explained the strong association between cotinine and the rs16969968/rs1051730 genotype. This suggests that rather than increased volume of smoke inhalation, there may be an alternative mechanism affecting the association between cotinine levels and rs16969968/rs1051730 variation.

It is possible that the mechanism underlying the association between the *CHRNA5-A3* genotype and cotinine levels is subject to plasticity across the lifespan. The genetic variation that characterizes the nAChR subunit may change its electrostatic balance and consequently its conformational state changes. We cannot exclude that such balance change may be subjected to modifications across the lifespan and manifest differently in young smokers and older smokers. Further *in vitro* and *in silico* studies are needed to explore this hypothesis.

## Acknowledgements

We are extremely grateful to all the families who took part in the ALSPAC study, the midwives for their help in recruiting them, and the whole ALSPAC team, which includes interviewers, computer and laboratory technicians, clerical workers, research scientists, volunteers, managers, receptionists and nurses. We are also very grateful to the volunteer researchers Sema Balaban and Boglarka Zilahi for the help to collect data at LSBU. The UK Medical Research Council and Wellcome Trust (Grant ref: 102215/2/13/2) and the University of Bristol provide core support for ALSPAC.

## Authors Contributions

TE, MRM, LD, and NJT conceived the work, GL conducted the studies, GL, MD, KD and LD collected the data, GL, VT, LM, JM analysed the data, SFO performed the modeling of α3α5β4 nAChR. GL, SR, JM, NJT, TE, MRM contributed to interpreting the data. GL, NJT and MRM drafted the manuscript and all authors reviewed and approved the final version of the manuscript.

This publication is the work of the authors and Glenda Lassi, Vanessa Tan, Liam Mahedy, Ana Sofia F. Oliveira, Maddy L. Dyer, Katie Drax, Lynne Dawkins, Stephen Rennard, James Matcham, Nicholas J. Timpson, Tim Eisen, Marcus R. Munafò, will serve as guarantors for the contents of this paper.

