## Supplementary material for "Exploration of the role of *CHRNA5-A3-B4* genotype in smoking behaviours"

**Structure Modeling of α3α5β4 nAChR**

Since the mechanism though which nAChR modulates these phenotypes is not perfectly understood, we generated a homology model of the human α3α5β4 nAChR in an attempt to clarify the structural and functional effect of the D398N mutation (Supplementary Figure 1). In this model, based on the templates of the human α4β2 (pdb code: 5kxi [1]) and the *Torpedo marmorata* (pdb code: 2bg9 [2]) nAChRs, D398 is located in the intracellular vestibule of the receptor. This locates D398 ~45 Å away from the “selectivity filter” (located at the intracellular end of the ion channel) and 114 Å away from the closest orthosteric binding site (Supplementary Figure 1A). MODELLER 9v17 [3] was used for generating the structure. The quality of the homology model was assessed by considering the restraint violations reported by MODELLER [3] combined with a Ramachandran analysis performed by the program PROCHECK [4]. The final model has 92.8% of the residues in the most favored regions and 7.1% in additional allowed regions.The homology model was inserted into a pre-equilibrated 1-palmitoyl-2-oleoyl-sn-glycero-3-phosphocholine (POPC) lipid bilayer using LAMBADA and InflateGRO2 [5] and, then, solvated using TIP3P water molecules [6]. The system was energy minimized with the steepest-descent

method in order to remove excessive strain. 10,000 steps of minimization with harmonic restraints applied to all non-hydrogen atoms were performed followed by further 10,000 steps restraining the C_α_ atoms only, ending with 10,000 steps with no restraints. The Amber ff99SB-ILDN [7] force field was used for the protein and the parameters for nicotine were taken from our previous work [8]. Additionally, the surface electrostatic distributions were determined for the wild-type and the D398N mutant using the APBS [9] software.

Additionally, we determined the electrostatic potential surface for the wild-type and the D398N mutant. The mutation of aspartate 398 to an asparagine affects the charge distribution of the intracellular vestibule region (Supplementary Figure 1B-C) due to the replacement of the negative charged residue by a neutral one. The intracellular domain is thought to play important roles in the stabilization and modulation of the receptor [10, 11] and it contributes to the charge selectivity of the ion channel [2]. In addition, previous studies have shown that nAChR activity changes according to the local electrostatic charge [12, 13]. Consequently, a mutation in this region is likely to affect the activity of the receptor.

The α5 nAChR subunit is an accessory subunit that does not contribute to the nicotine binding site [14-16] and the location of D398 is too far to directly affect the sensitivity of agonistic binding. In fact, D398 is located in the intracellular vestibule of the receptor; the intracellular domain is highly variable among the subunits but hosts sites responsible for modifications and interactions with cytoplasmatic elements [17]. Furthermore, the intracellular vestibule has an overall negative charge that enhances the catonic selectivity and contributes to the high conductance of the nAChR channel. Thereby the D398N mutation, by substituting a negatively charged residue with a neutral one, is likely to alter the cations trafficking. Our model confirms the findings by Frahm and colleagues [13] who built a model exclusively based on the *Torpedo marmorata* as the sequence and the structure of the human α4β2 was not available yet.

The kinetic rate at which nAChRs change conformational states (i.e., rest, open, and desensitized) depends on many factors including the subunit structure [17]. It is plausible that, since nicotine inhaled though cigarette smoke activates and desensitizes nAChRs, a mutation (a substitution of an amino acid that affects the nAChR subunit) that changes the electrostatic balance of the receptor may have a role in the response to nicotine. Nevertheless, further experiments are needed to explore this hypothesis.

**Supplementary Figure 1. Model of the human α3α5β4 nAChR**.

The α3, α5 and β4 subunits are colored in yellow, cyan and green, respectively. The orange sticks represent nicotine (which bind to the orthosteric binding sites). The position of D398 is highlighted with red spheres (A). Electrostatic potential surface representations for the wild-type (B) and for the D398N mutant α3α5β4 nAChR (C). The potential varies between -5 and +5 kT/e, as shown in the colors bar, with red and blue representing negative and positive potentials, respectively.

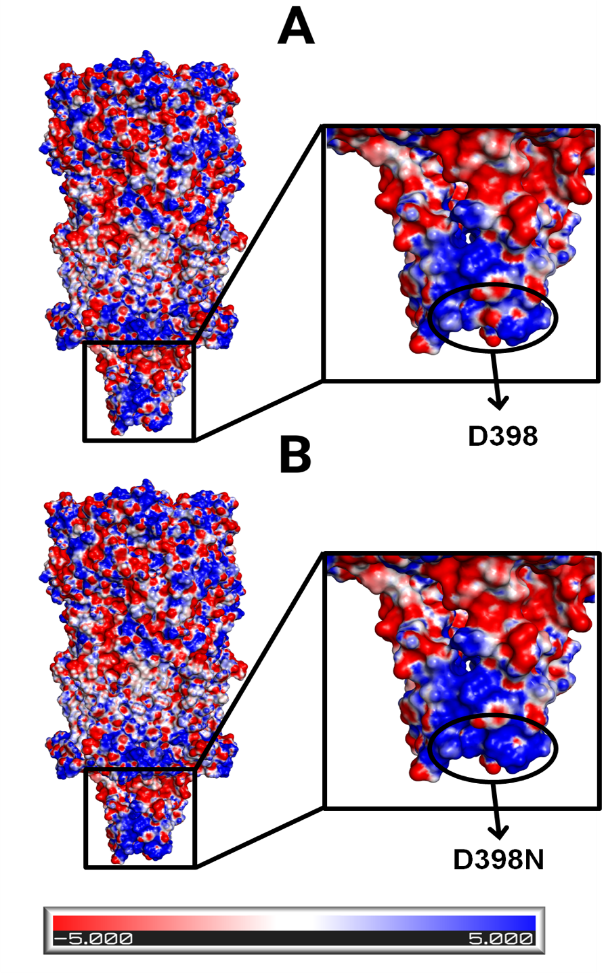

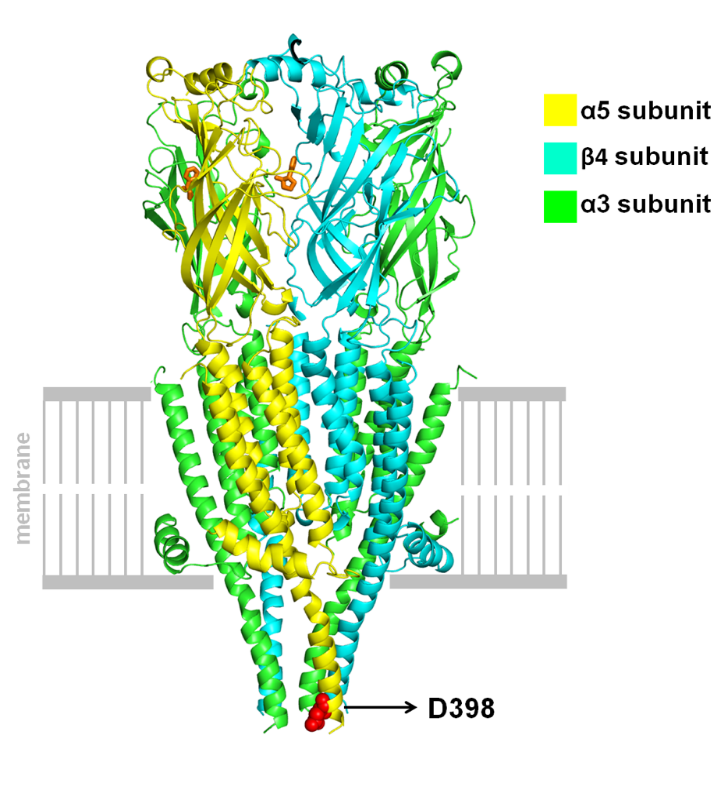

**A**

**B**

**C**

**Genotyping.** ALSPAC participants’ genotype was known a priori although the researcher was blind to the genotypes of all participants. For non ALSPAC participants, DNA samples were obtained using the Oragene DNA oral collection kit OG-500 (DNA Genotek Inc., Canada) to collect and store saliva. The samples were genotyped for rs1051730 by KASP genotyping assay (LCG Genomics), using the following SNP sequence:

TTTCCAGGTGGACGACAAGACCAAAGCCTTACTCAAGTACACTGGGGAGGTGACTTGGATACCTCCGGCCATCTTTAAGAGCTCCTGTAAAATCGACGTGACCTACTTCCCGTTTGATTACCAAAACTGTACCATGAAGTTCGGTTCCTGGTCCTACGATAAGGCGAAAATCGATCTGGTCCTGATCGGCTCTTCCATGAACCTCAAGGACTATTGGGAGAGCGGCGAGTGGGCCATCATCAAAGCCCCAGGCTA[C/T]AAACACGACATCAAGTACAACTGCTGCGAGGAGATCTACCCCGACATCACATACTCGCTGTACATCCGGCGCCTGCCCTTGTTCTACACCATCAACCTCATCATCCCCTGCCTGCTCATCTCCTTCCTCACTGTGCTCGTCTTCTACCTGCCCTCCGACTGCGGTGAGAAGGTGACCCTGTGCATTTCTGTCCTCCTCTCCCTGACGGTGTTTCTCCTGGTGATCACTGAGACCATCCCTTCCACCTCGCTGGTC.

**Supplementary Table 1. Nicotine challenge study: associations between genotype and possible covariates.**

| **Confounders** | Ethnicity | N | Coefficient | 95% CI | P-value |
| --- | --- | --- | --- | --- | --- |
| **Sex** | All | 520 | OR -0.05 | -0.34 to 0.23 | 0.700 |
|  | European | 400 | OR -0.06 | -0.37 to 0.26 | 0.731 |
| **Age** | All | 520 | Beta coeff. -0.14 | -0.73 to 0.45 | 0.643 |
|  | European | 400 | Beta coeff. -0.05 | -0.60 to 0.71 | 0.876 |

OR = odds ratio.

**Supplementary Table 2.** **Nicotine challenge study: values of BP, HR, VAS and PANAS according to research site.**

|  | **UoB** | | | **LSBU** | | |
| --- | --- | --- | --- | --- | --- | --- |
|  | **Mean (SD)**  **Range** | | | | | |
|  | **Baseline** | **Change score at 5 min** | **Change score at 10 min** | **Baseline** | **Change score at 5 min** | **Change score at 10 min** |
| **BP sys** | 114.90 (13.50)  83 – 172 | 4.06 (10.02)  -68 – 36 | 0.51 (10.12)  -78 – 37 | 117.3 (12.95)  92 – 160 | 3.35 (12.14)  -54 – 24 | 0.49 (12.72)  -50 – 43 |
| **BP dia** | 74.61 (9.05)  53 – 128 | 7.09 (9.35)  -59 – 51 | 3.66 (8.49)  -53 – 43 | 70.35 (12.42)  49 – 140 | 4.55 (13.60)  -80 – 33 | 3.30 (14.96)  -80 – 52 |
| **HR** | 74.87 (14.76)  41 – 127 | 14.66 (9.99)  -29 – 70 | 9.88 (8.55)  -32 – 38 | 75.93 (11.83)  50 – 101 | 13.96 (8.83)  -8 – 34 | 9.54 (8.69)  -14 – 30 |
| **VAS** | 5.54 (7.44)  0 – 55 | 25.76 (15.86)  -12 – 79 | 16.93 (15.05)  -6 – 86 | 3.67 (4.25)  0 – 20 | 24.44 (19.16)  0 – 86 | 15.35 (15.77)  -3 – 62 |
| **PAs** | 23.54 (6.90)  7 – 44 |  | -5.06 (5.84)  -27 – 27 | 27.65 (9.88)  12 – 74 |  | -5.38 (8.41)  -48 – 8 |
| **NAs** | 11.91 (2.92)  1 – 26 |  | 2.26 (4.13)  -12 – 20 | 12.24 (3.06)  9 – 26 |  | 0.97 (4.39)  -6 – 24 |

Mean, standard deviation (SD) and range of all outcomes (baseline and change score at 5 and 10 minutes) according to research site (UoB = University of Bristol; LSBU = London South Bank University).

**Supplementary Table 3.** **Smoking topography study: associations between genotype and possible confounders.**

| **Confounders** | N | Beta coefficient | 95% CI | P-value |
| --- | --- | --- | --- | --- |
| **Sex** | 100 | -0.26 | -0.16 to 0.11 | 0.706 |
| **CPD** | 100 | 0.46 | -0.88 to 1.80 | 0.500 |

CPD = cigarettes per day.

**Supplementary Table 4. Smoking topography study: Characteristics of daily smokers**

|  |  | **ALL**  **N = 123** | | **YOUNG ADULTS**  **N = 100** | | **ADULTS**  **N = 15** | | **OLDER ADULTS**  **N = 8** | |
| --- | --- | --- | --- | --- | --- | --- | --- | --- | --- |
| **Variable** |  | **N** | **Mean (SD) Range** | **N** | **Mean (SD) Range** | **N** | **Mean (SD)**  **Range** | **N** | **Mean (SD) Range** |
| **Age (years)** |  | 123 | 24.74 (8.89)  18 – 67 | 100 | 21.64 (1.51)  18 – 24 | 14 | 29.13 (4.53)  25 – 40 | 8 | 55.25 (7.92)  47 – 67 |
| **Cotinine levels** |  | 121 | 145.05 (107.38)  0.2 – 514.1 | 98 | 131.13 (96.33)  0.2 – 502.99 | 15 | 159.54 (97.34)  25.4 – 392. 5 | 8 | 288.4 (153.17)  110.9 – 514.1 |
| **FTND score (1-10)** | <4 | 75 | 1.69 (1.11)  0 – 3 | 66 | 1.62 (1.11)  0 – 3 | 6 | 1.83 (1.17)  0 – 3 | 3 | 3.00 (0.00)  3 – 3 |
|  | ≥4 | 48 | 5.12 (1.30)  4 – 9 | 34 | 5.09 (1.33)  4 – 9 | 9 | 5.11 (1.45)  4 – 7 | 5 | 5.40 (0.89)  4 – 6 |
| **CPD self-report** | ≤ 10 | 74 | 6.91 (2.28)  1 – 10 | 64 | 6.78 (2.30)  1 – 10 | 9 | 7.50 (2.03)  4 – 10 | 1 | 10.00 (0.00)  10 – 10 |
|  | >10 | 49 | 15.68 (4.14)  11 – 30 | 36 | 15.12 (3.73)  11 – 30 | 6 | 14.50 (2.07)  12 – 17 | 8 | 19.60 (5.59)  12 – 25 |

Age, FTND and CPD mean values, standard deviation (SD) and range of values for all genotyped participants and according to age group. FTND = Fagerström Test of Nicotine Dependence; CPD = cigarettes per day; YP = young people.

**Supplementary Table 5. Smoking topography study: cardiovascular, craving and mood.**

| **OUTCOME** | | **YOUNG PEOPLE** | | **Paired two tailed t-test** |
| --- | --- | --- | --- | --- |
|  |  | **N** | **Mean (SD) Range** |  |
| **HR** | **PRE** | 96 | 76.62 (11.40) 47-106 | t _(95)_ = -13.17  P-value < 0.001 |
|  | **POST** | 96 | 88.47 (12.43) 55-121 |  |
| **BP systolic** | **PRE** | 98 | 115.57 (15.27) 91-176 | t _(97)_ = -3.71  P-value < 0.001 |
|  | **POST** | 98 | 119.98 (13.56) 87-150 |  |
| **BP diastolic** | **PRE** | 98 | 72.44 (9.12) 48-110 | t _(97)_ = -5.45  P-value < 0.001 |
|  | **POST** | 98 | 76.29 (8.87) 57-97 |  |
| **QSU** | **PRE** | 100 | 28.86 (10.86) 10-58 | t _(99)_ = 16.08  P-value < 0.001 |
|  | **POST** | 100 | 14.33 (4.96) 10-33 |  |
| **Positive Affect** | **PRE** | 100 | 26.25 (6.04) 11-43 | t _(99)_ = 3.33  P-value = 0.006 |
|  | **POST** | 100 | 24.71 (6.97) 11-43 |  |
| **Negative Affect** | **PRE** | 100 | 12.08 (2.93) 10-24 | t _(99)_ = 0.96  P-value = 0.17 |
|  | **POST** | 100 | 11.08 (2.78) 10-24 |  |

Mean, standard deviation and range of values pre and post smoking a cigarette in the laboratory. T-tests were performed to assess whether smoking affected heart rate (HR), systolic and diastolic blood pressure (BP), craving measured with the Questionnaire of Smoking Urges (QSU) and mood assessed with the Positive and Negative Affect Scale (PANAS).

**Supplementary Table 6. Smoking topography study: summary statistics of the smoking topography outcomes.**

| **SMOKING TOPOGRAPHY OUTCOMES** | | **N** | **Mean (SD) Range** |
| --- | --- | --- | --- |
| **LAB - 1 cigarette** | **Volume (ml)** | 92 | 1241.01 (887.35)  208.8-3928.9 |
| **LAB – 1 puff** | **Mean volume (ml)** | 92 | 58.59 (36.40)  15.26-218.2 |
| **HOME - 1 cigarette** | **Mean volume (ml)** | 94 | 927.92 (463.47)  248.38-2338 |
| **HOME – 1 puff** | **Mean volume (ml)** | 94 | 51.72 (26.12)  14.40-155.90 |
| **HOME – 24 hour** | **Total volume inhaled (ml)** | 92 | 78020.88 (6125.10)  993.90-37333.40 |

Mean, standard deviation and range of values.

**Supplementary Table 7.** **Smoking topography study: correlations between smoking topography outcomes measured in the laboratory and measured at home.**

| **VARIABLES** | **r** | **p-value** |
| --- | --- | --- |
| 1 cigarette volume | 0.72 | <0.0001 |
| 1 puff volume | 0.78 | <0.0001 |
| CPD | 0.73 | <0.0001 |

CPD = cigarettes smoked per day (which was self-reported during the laboratory session and measured by the smoking device in the following 24h)

**Supplementary Table 8. Smoking topography study: summary statistics of additional smoking topography outcomes.**

| **SMOKING TOPOGRAPHY OUTCOMES** | | **N** | **Mean (SD) Range** |
| --- | --- | --- | --- |
| **1 cigarette** | **Duration (s)** | 86 | 269.82 (85.95)  122.55-51.06 |
| **1 puff** | **Duration (s)** | 90 | 2.08 (0.77)  0.86-4.85 |
| **Inter-puff interval** | **Duration (s)** | 86 | 12.93 (5.45)  122.55-510.57 |
| **Puffs** | **Total number** | 93 | 21.29 (8.93)  8-51 |

Mean, standard deviation and range of values.

**Supplementary Table 9. Smoking topography study: association between genotypes and further smoking topography outcomes.**

| **GENOTYPE** | **Model** | **N** | **Beta coeff** | **95% CI** | **P-value** |
| --- | --- | --- | --- | --- | --- |
| **Duration of 1 cigarette (s)** | 1 | 86 | -0.07 | -0.16 to 0.01 | 0.105 |
|  | 2 | 86 | -0.07 | -0.16 to 0.02 | 0.135 |
|  | 3 | 86 | -0.05 | -0.13 to 0.04 | 0.285 |
| **Mean duration of puffs (s)** | 1 | 90 | 0.04 | -0.06 to 0.13 | 0.470 |
|  | 2 | 90 | 0.03 | -0.07 to 0.13 | 0.520 |
|  | 3 | 90 | 0.01 | -0.08 to 0.11 | 0.806 |
| **Inter puff interval (s)** | 1 | 86 | -0.05 | -0.15 to 0.06 | 0.382 |
|  | 2 | 86 | -0.05 | -0.16 to 0.05 | 0.318 |
|  | 3 | 86 | -0.04 | -0.14 to 0.07 | 0.458 |
| **Total number of puffs** | 1 | 93 | -0.07 | -0.18 to 0.03 | 0.174 |
|  | 2 | 93 | -0.07 | -0.18 to 0.04 | 0.202 |
|  | 3 | 93 | -0.06 | -0.17 to 0.04 | 0.242 |

Regressions were 1. Unadjusted; 2. Adjusted for sex; 3. Adjusted sex and cigarettes smoked per day. Genotype was tested for associations with additional measures collected in the laboratory (total number of puffs per cigarette and all the temporal variables).

**Supplementary Table 10. Smoking topography study: latent profile analysis.**

|  | #param | AIC | BIC | SBIC | Entropy | Min class | LRT | BLRT |
| --- | --- | --- | --- | --- | --- | --- | --- | --- |
| 1 class | 8 | 3387 | 3411 | 3385 | - | - | - | - |
| 2 class | 13 | 3293 | 3331 | 3290 | 0.79 | 30.6% | 0.036 | 0.041 |
| 3 class | 18 | 3269 | 3322 | 3265 | 0.80 | 11.0% | 0.112 | 0.118 |

Comparison of model fit indices comparing 1 to 3 profiles. #param: number of parameters; AIC: Akaike information criterion; BIC: Bayesian information criteria; SBIC: sample-size-adjusted Bayesian information criterion LRT: Lo-Mendell-Rubin likelihood ratio test; BLRT: bootstrap likelihood ratio test.
